# Mitochondrial subpopulations in oocytes and cumulus cells exhibit distinct age-associated changes and selective plasticity in response to NMN supplementation

**DOI:** 10.64898/2026.03.25.714214

**Authors:** Andrew J. Piasecki, Hannah Sheehan, Paula Ledo Hopgood, Jonathan L. Tilly, Dori C. Woods

**Affiliations:** Department of Biology, Northeastern University, Boston, MA, United States; SauveBio Inc., Worcester, MA, United States; University of Miami Miller School of Medicine, Miami, FL, United States

**Keywords:** mitochondrial heterogeneity, oocyte quality, cumulus granulosa cells, NMN, reproductive aging, membrane potential, mtDNA

## Abstract

**Background:** Mitochondrial dysfunction is a leading contributor to the decline in oocyte quality associated with maternal aging. Prior investigations of mitochondrial function in the ovarian follicle have largely treated the mitochondrial pool as a homogeneous population, reporting aggregate values that may obscure biologically meaningful differences between distinct mitochondrial subpopulations. The present study addresses this limitation by characterizing mitochondrial subpopulation dynamics in oocytes and cumulus granulosa cells at single-organelle resolution using fluorescence-activated mitochondria sorting (FAMS).

**Results:** Analysis of the aggregate mitochondrial population in mouse oocytes revealed no significant age-associated differences in mitochondrial DNA copy number or membrane potential, a result that would previously have been interpreted as evidence of minimal age-related mitochondrial change. Subpopulation analysis revealed this conclusion to be incomplete: aged oocytes showed significantly elevated mitochondrial DNA copy number specifically within the high membrane potential and small mitochondrial subpopulations, with no significant differences in the low membrane potential or large subpopulations. NMN supplementation normalized mitochondrial DNA copy number in the high membrane potential and small subpopulations toward young levels while producing an opposing effect in large mitochondria, demonstrating subpopulation-specific rather than uniform rejuvenation. In cumulus cells, significant age-associated changes were detectable at the aggregate level, including a reduction in mitochondrial DNA copy number and an elevation in membrane potential, and subpopulation analysis further resolved these findings. The age-associated reduction in cumulus cell mitochondrial DNA copy number was driven predominantly by the high membrane potential subpopulation. NMN supplementation exerted opposing effects on small and large cumulus cell mitochondrial subpopulations, increasing mitochondrial DNA copy number above both young and aged levels in small mitochondria while further reducing it below aged levels in large mitochondria.

**Conclusions:** Viewing the mitochondrial pool as a heterogeneous mixture of functionally distinct subpopulations rather than a uniform population reveals age-associated alterations in oocytes and cumulus cells that are undetectable by aggregate analysis. NMN supplementation exerts subpopulation-specific effects in both cell types, identifying specific mitochondrial subtypes as more precise targets for future mechanistic investigation of age-associated infertility than the mitochondrial pool considered in aggregate.

## Introduction

The ovary is one of the first organs to fail with age, well out of sync with the chronological lifespan of women. This decline in ovarian function is tightly associated with a decline in both the quantity and quality of oocytes available for fertilization [1,2]. Of the many mechanisms proposed as being responsible for driving deterioration of egg quality with age, mitochondrial dysfunction appears central to the process [3]. Prior studies have shown that there is a marked decrease in mitochondrial number and function in oocytes with age, associated with abnormal mitochondrial distribution patterns [4–9]. Importantly, the mitochondrial role in ovarian function with age is not restricted to the oocyte, as mitochondria play a major role in other aspects of ovarian function including steroid hormone biosynthesis in follicular somatic cells [10], and even ovarian stromal cells have been reported to demonstrate compromised mitochondrial bioenergetics with age [11].

It is important to note that mitochondria are not a homogeneous pool of organelles; they display a wide degree of diversity not only between cell types, but even within individual cells. Mitochondria have been shown to vary in both form, including size, ultrastructure, cristae density, physical networking, and mitochondrial DNA (mtDNA) content, and function, including membrane potential (ΔΨ_M_), apoptotic priming, hormone biosynthesis, and modulation of nuclear gene expression through retrograde signaling [12–14]. This variation can be studied by classifying mitochondria into subpopulations on the basis of these characteristics. While previous studies lacked the technology to discern these nuances on an intracellular level, the development of fluorescence-activated mitochondria sorting (FAMS) allows for the identification of mitochondrial characteristics on a single-organelle basis [13,14,15]. This increased resolution provides a means to define clear subpopulations of mitochondria on the basis of physical and functional differences. Given the direct link between a mitochondrion’s physical characteristics and its role within the cell, defining the mitochondrial subpopulations present within the ovarian follicle and evaluating how aging affects their dynamics is important to fully understand the impact of mitochondrial function on female fertility.

Recent studies have shown that age-associated infertility can be lessened by nicotinamide mononucleotide (NMN), first reported by Bertoldo et al. to rejuvenate oocyte quality in aged mice [16]. Continued research has since shown NMN supplementation to improve both oocyte quality and quantity in aged individuals [17–19], as well as conferring protective effects against conditions such as obesity, heat stress, and chemical insult [20–22]. NMN supplementation also directly impacts mitochondria within the ovary, with treatment of aged individuals producing effects including improved mitochondrial distribution within oocytes, improved hormone production by granulosa cells, reduced reactive oxygen species levels, increased cristae density, and improved mitophagy and mitochondrial biogenesis [9,23,24,18]. Given that mitochondrial dysfunction is a leading cause of age-associated infertility, the improved mitochondrial function that results from NMN supplementation is likely to contribute to its restorative effects on overall ovarian function. However, prior investigation of NMN supplementation on mitochondrial function in the context of age-associated infertility has been conducted at the whole-cell level, leaving the subpopulation-specific effects of NMN on individual mitochondrial subtypes uncharacterized. Beyond its potential as a fertility intervention, NMN supplementation represents a useful pharmacological tool for probing the plasticity of the aged mitochondrial pool, since its ability to alter mitochondrial characteristics in aged individuals provides a means to assess whether age-associated changes are reversible or fixed.

Prior work in non-reproductive cell types has shown that aging increases mitochondrial heterogeneity in a subpopulation-specific manner, with distinct mitochondrial subpopulations within the same cell exhibiting markedly different membrane potential profiles in aged versus young individuals [31]. In the context of the female germline specifically, it has been proposed that the mitochondrial anomalies associated with oocyte aging may be reflective of changes in specific mitochondrial subpopulations rather than uniform alterations across the entire organelle pool [3], though this hypothesis had not been directly tested prior to the present study. Based on this prior evidence, together with the established heterogeneity of mitochondria within individual cells with respect to size, ΔΨ_M_, and mtDNA content [12–15], we hypothesized that aging differentially affects distinct mitochondrial subpopulations in oocytes and cumulus granulosa cells, and that these subpopulation-specific changes would be obscured by aggregate analysis. To test this hypothesis, FAMS was employed to analyze mitochondria from young, aged, and NMN-supplemented aged mice at single-organelle resolution, with NMN supplementation used as a pharmacological probe of mitochondrial plasticity, with the aim of identifying age-associated alterations in mitochondrial subpopulation dynamics and determining the capacity of NMN supplementation to restore these subpopulations toward the mitochondrial profile observed in young individuals.

## Materials and Methods

### Animals and Reagents

Young C57BL/6 female mice (2 months of age upon arrival) were obtained from the Jackson Laboratory (Bar Harbor, ME). Aged C57Bl/6 female mice (12 months of age upon arrival) were obtained from the National Institute of Aging (Bethesda, MD). All experiments described herein were reviewed and approved by the Institutional Animal Care and Use Committee at Northeastern University.

### Nicotinamide Adenine Dinucleotide (NAD+) Repletion

For NAD+ repletion, 0.5 g/L of nicotinamide mononucleotide (NMN) was added as a supplement to the water supply of 12-month-old female C57BL/6 mice for 28 days (4 weeks). The NMN-supplemented water supply was replenished as-needed, and at minimum every two weeks. After the 28-day treatment, mice were given hormone injections for superovulation to collect cumulus-oocyte-complexes (COCs) for downstream analysis.

### Collection of Cumulus-Oocyte-Complexes

Ovulation was induced by intraperitoneal injection of pregnant mare serum gonadotropin (PMSG; 5 IU for 2-month-old mice and 10 IU for 12-month-old mice; ProspecBio) followed by human chorionic gonadotropin (hCG; 5 IU for 2-month-old mice and 10 IU for 12-month-old mice; ProspecBio) 46-48 h later. Mice were sacrificed by cervical dislocation 13-14 h post-hCG injection, and their oviducts were collected by dissection. COCs were released from the oviducts as an intact clutch by gently tearing the ampulla with an insulin syringe.

### Isolation of Cumulus Cells and Oocytes

For experiments where cumulus cells and oocytes were analyzed independently, COCs were incubated for 2 min in 80 IU/mL of hyaluronidase (Sigma Aldrich) at 37°C to separate the cumulus cells from the oocytes. After the removal of all oocytes by mouth pipette, the dish, which then contained only cumulus cells, was flooded with 1.5 mL of PBS and transferred to a microcentrifuge tube. The cumulus cell suspension was then centrifuged at 500 g for 5 min. Denuded oocytes were washed three times with HEPES-buffered M2 medium (Sigma Aldrich) supplemented with 0.4% BSA at 37°C.

### MitoTracker Green FM and JC-1 Labeling for Fluorescence-activated Mitochondria Sorting (FAMS)

For intracellular mitochondrial samples, oocytes and cumulus cells were either transferred to 200 µL of ice-cold lysis buffer (10 mM NaCl, 1.5 mM MgCl₂, 10 mM Tris-HCl pH 7.6, 1X Halt Protease Inhibitor Cocktail in PBS) by a glass capillary tube (individual oocytes) or brought to a single cell suspension in 200 µL ice-cold lysis buffer (cumulus cells). To lyse the cells, samples were vortexed for 1 min and centrifuged at 12,000 g for 5 min at 4°C. After removal of supernatant, all samples were resuspended in PEB buffer [0.5% BSA, 2 mM EDTA in PBS; [13]] and stained with 100 nM MitoTracker Green (MTG; Invitrogen) and 1 µM JC-1 (Marker Gene Technologies) for 15 min at room temperature, protected from light. Samples were washed with PEB buffer, centrifuged at 12,000 g, and resuspended in cold sheath fluid before transferring to polystyrene FACS tubes for FAMS analysis and sorting.

### Calibration of Flow Cytometer for FAMS

All analyses were completed using a special-order research product BD FACS Aria III, fitted with a PMT detector for forward light scatter (FSC) of a 488 nm laser, allowing for an increased dynamic range of small particle detection over standard photodiode detectors. To achieve high-resolution detection of subcellular-sized calibration beads (<1 μm), the instrument detection threshold was routinely set to side light scatter (SSC) 200. Following standard instrument calibration of laser area scaling and delay, a mixture of size calibration beads (Spherotech and Life Technologies) was run to optimize light scatter voltages for visualization of events between 0.22 and 4 μm. For all analyses, at least 3 × 10⁴ events were analyzed per sample. Data were acquired using BD FACSDiva software (version 8.0.2) and then further analyzed using FlowJo (v10.8.1) and Microsoft Excel (v16.61.1).

### Determination of Mitochondrial DNA (mtDNA) Copy Number

To determine mtDNA copy number on a per-organelle basis, individual mitochondria were sorted from oocyte and cumulus cell samples into 96-well PCR plates containing 1 µL of mitochondrial lysis buffer (10 mM EDTA, 0.5% SDS, 0.1 mg/mL Proteinase K in qPCR grade water). After the sorting of one mitochondrion into each well of the 96-well plate was complete, all wells were immediately overlaid with mineral oil to prevent evaporation and were incubated at 37°C for 1 h before being transferred to a -80°C freezer until preparation for single molecule polymerase chain reaction (smPCR). Lysed samples were diluted across 32 PCR wells to prevent inhibitory effects of mitochondrial lysis buffer on amplification, and to ensure templates were individually segregated into separate wells. smPCR was performed using Ex Taq DNA Polymerase, Hot Start Version (Takara) with LA PCR buffer (Takara) to generate 331 base pair amplicons. For primer information, please see MacDonald et al. 2019 [13]. Following gel electrophoresis, mtDNA copy number was determined by manually counting the number of bands present in each 32-well sample, since the smPCR method has been optimized in such a way that each band represents one mtDNA copy.

### Statistical Analysis

Analysis of statistical significance for mtDNA copy number and JC-1-based ΔΨ_M_ data was conducted using GraphPad Prism (v9.3.1) with a threshold for statistical significance of P<0.05. The data generated using the young, aged, and NAD+ repleted sample groups regarding mtDNA copy number and ΔΨ_M_ were, in most cases, non-normally distributed (according to the Anderson-Darling test, the D-Agostino & Pearson test, the Shapiro-Wilk test, and the Kolmogorov-Smirnov test). Therefore, the non-parametric Mann-Whitney U two-tailed test was used to determine statistical significance between sample groups. For ΔΨ_M_ analysis, a sample size of 6 biological replicates (12-month age group) was the minimum used to determine statistical significance for each experimental group, with the largest sample size being 12 biological replicates (2-month age group). For mtDNA copy number determination, the minimum sample size was 24 mitochondria. No sample-size calculation was performed, however, all attempts at replication were successful.

## Results

### Mitochondrial subpopulations are present in both oocytes and cumulus cells

To establish whether mitochondrial subpopulations are present within oocytes and cumulus cells, FAMS was employed to analyze the size, ΔΨ_M_, and mtDNA copy number of individual organelles. Typical size gates for the sorting of mouse oocyte and cumulus cell mitochondria were drawn to encompass size calibration beads ranging in size from 0.22 to 2 μm (Figure 1), consistent with transmission electron microscopy-based evaluation of oocyte mitochondrial size range in situ [13,30]. Mitochondria were identified by size and by positive fluorescence using the mitochondrial-specific dye MTG. For the MTG gate, an unstained sample was used to establish a negative population with respect to MTG (FITC)-positive events, and a gate was drawn to exclude all but up to ∼0.5% of autofluorescent events to correct for experimental variation (Figure 2). Compensation was not applied to samples labeled with JC-1, which measures ΔΨ_M_, due to the emission properties of the dye.

**Figure 1.**
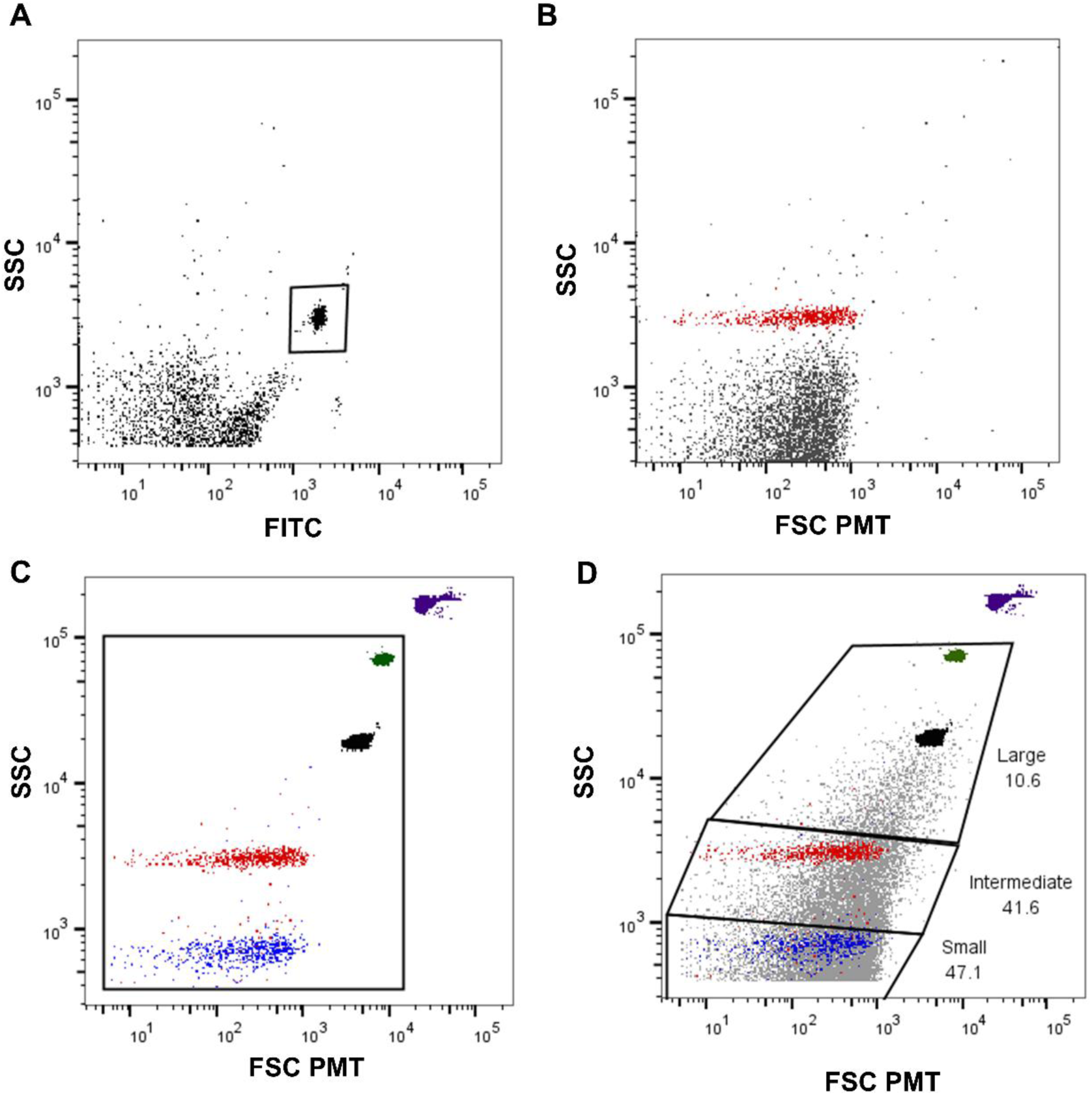
Experimental workflow and fluorescence-activated mitochondria sorting (FAMS) gating strategy. Cumulus-oocyte complexes (COCs) were collected from young (2-month-old), aged (12-month-old), and NMN-supplemented aged C57BL/6 female mice. Following hyaluronidase treatment and mechanical separation, oocytes and cumulus granulosa cells were isolated independently and subjected to cell lysis to release mitochondria into suspension. Each sample was analyzed separately by FAMS. Sequential gating was applied to identify and define mitochondrial subpopulations: an initial size gate (0.22–2 μm) was applied to FSC PMT vs. SSC plots to isolate events within the mitochondrial size range from cellular debris (left plot); MitoTracker Green FM (MTG) fluorescence in the FITC channel was then used to confirm mitochondrial identity within the size-gated population (center plot); size-gated, MTG-positive events were subsequently divided into small (<0.3 μm), intermediate (0.31–0.6 μm), and large (>0.6 μm) subpopulations based on FSC PMT values (right plot). Each sample was processed and analyzed independently.

**Figure 2.**
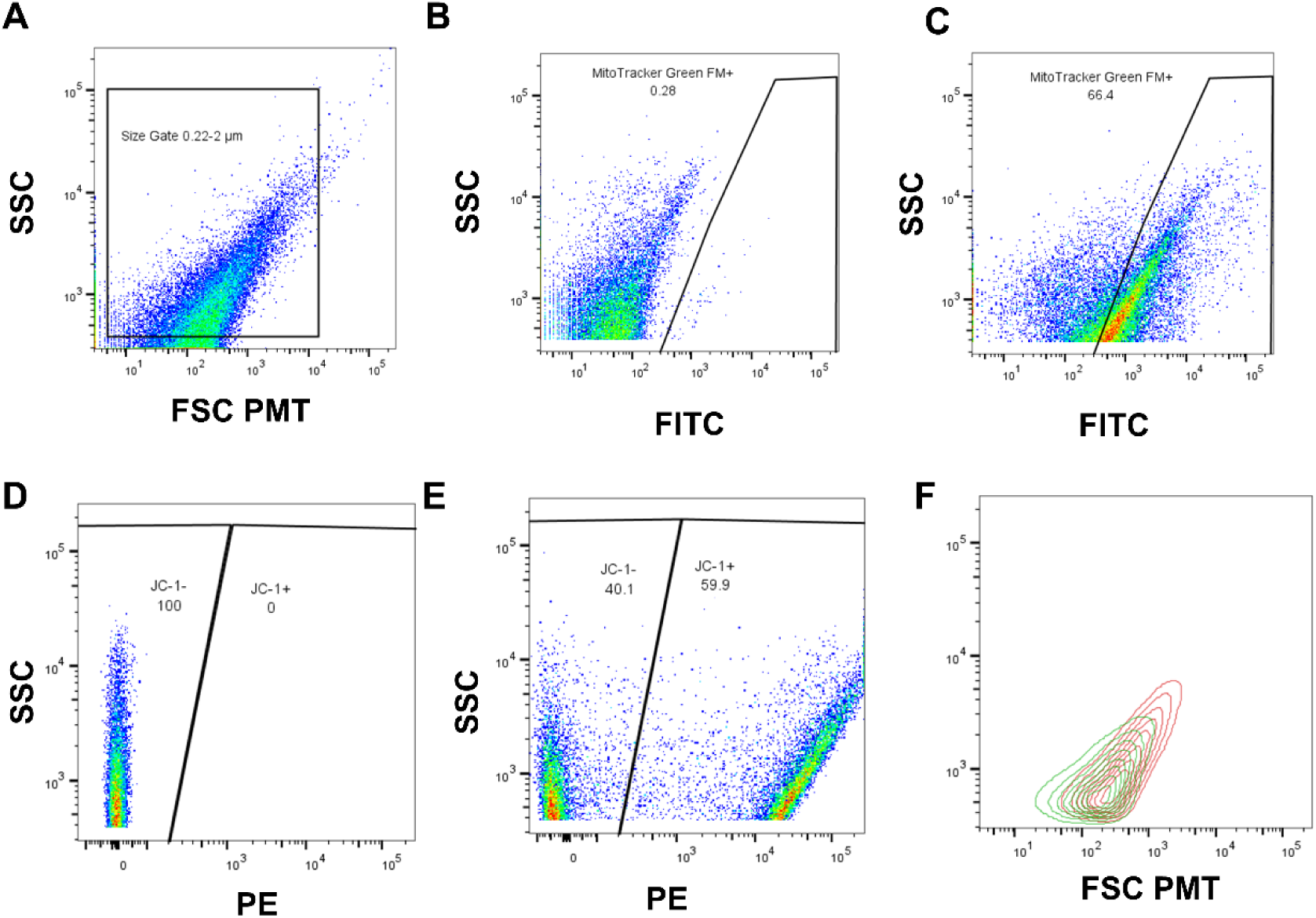
Representative FAMS gating strategy for assessment of mitochondrial membrane potential (ΔΨM). (A) Size-gated events from a representative mitochondrial sample shown on FSC PMT vs. SSC axes. The size gate (0.22–2 μm) is indicated. (B) Negative control sample stained with MTG only, used to establish the MTG-negative baseline and set the MTG gate threshold. Events falling above the gate line (MTG+) represent fewer than 0.5% of autofluorescent events and define the false-positive threshold. (C) Representative MTG-stained sample showing MTG-positive mitochondria above the gate threshold. (D) JC-1 staining of an MTG-only sample (no JC-1) used to establish JC-1 gating in the PE channel. MTG-positive events are gated as JC-1-negative (ΔΨM-negative, left gate) and JC-1-positive (ΔΨM-positive, right gate). (E) Representative JC-1-stained sample showing the distribution of ΔΨM-negative (MTG+JC-1−) and ΔΨM-positive (MTG+JC-1+) mitochondrial populations. (F) Size overlay of ΔΨM-positive (high ΔΨM) and ΔΨM-negative (low ΔΨM) mitochondrial subpopulations shown as contour plots on FSC PMT vs. SSC axes, demonstrating the size heterogeneity present within each membrane potential subpopulation. Compensation was not applied to JC-1-stained samples due to the emission properties of the dye. All analyses were performed using a minimum of 3 × 10⁴ events per sample.

Variations in size, ΔΨ_M_, and mtDNA copy number were identified in both oocytes and cumulus cells, confirming the presence of distinct mitochondrial subpopulations in both cell types. The identified subpopulations were present in both young and aged samples, though their relative frequencies varied significantly in specific comparisons detailed below.

### Oocyte mitochondria mtDNA copy number varies significantly with age in specific mitochondrial subpopulations

Oocyte mitochondria were assessed on the basis of size, ΔΨ_M_, and mtDNA copy number through the use of FAMS and smPCR. Oocytes were isolated from COCs collected from young (2-month-old) and aged (12-month-old) mice, as well as from aged mice supplemented with NMN. Analysis of the aggregate oocyte mitochondrial population showed that age had no significant impact on mtDNA copy number (Figure 3A; mean = 0.732 copies per mitochondrion in young [n = 56, SEM = 0.249], 1.484 copies per mitochondrion in aged [n = 64, SEM = 0.527]) or ΔΨ_M_ (Figure 3B; 41.60% ΔΨM-positive in young [n = 12, SEM = 6.092], 39.40% in aged [n = 8, SEM = 10.360, p > 0.999]). NMN supplementation did not significantly impact aggregate ΔΨ_M_ relative to either age group [n = 8, SEM = 0.796], and the average aggregate mtDNA copy number in NMN-supplemented aged mice (0.147 copies per mitochondrion) was significantly reduced compared to untreated aged mice but was not significantly different from young mice [n = 34, SEM = 0.062]. These aggregate findings, considered alone, would suggest that oocyte mitochondria are not substantially altered by age or NMN supplementation. However, subpopulation analysis reveals a more complex picture.

**Figure 3.**
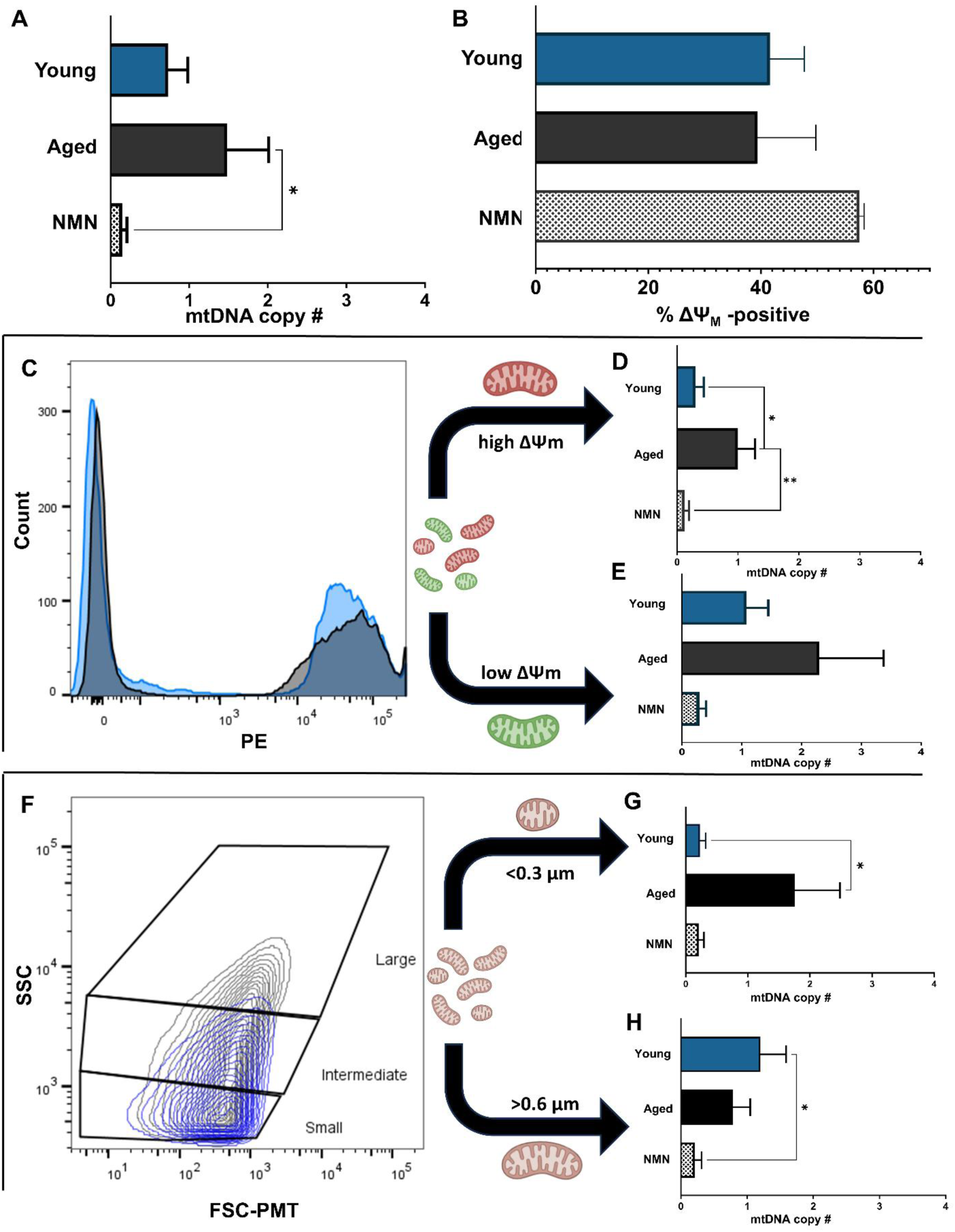
Age- and NMN-associated changes in mtDNA copy number in oocyte mitochondrial subpopulations. Mitochondria isolated from oocytes of young (2-month-old, blue), aged (12-month-old, black), and NMN-supplemented aged (hatched gray) C57BL/6 female mice were analyzed by FAMS and assessed for mtDNA copy number by single-molecule PCR (smPCR). (A) Mean mtDNA copy number per mitochondrion across the total oocyte mitochondrial population. (B) Percentage of ΔΨM-positive mitochondria across the total oocyte mitochondrial population. (C) Representative JC-1 PE channel histogram showing the distribution of ΔΨM-positive (high ΔΨM, right peak) and ΔΨM-negative (low ΔΨM, left peak) mitochondria in young (light blue) and aged (dark gray) oocyte samples. Curved arrows indicate separation of mitochondria into high and low ΔΨM subpopulations for downstream mtDNA analysis. (D) Mean mtDNA copy number in the high ΔΨM oocyte mitochondrial subpopulation. (E) Mean mtDNA copy number in the low ΔΨM oocyte mitochondrial subpopulation. (F) Representative FSC PMT vs. SSC contour plot showing the size distribution of oocyte mitochondria with small (<0.3 μm), intermediate (0.31–0.6 μm), and large (>0.6 μm) size gates indicated. Curved arrows indicate separation into small and large subpopulations for downstream mtDNA analysis. (G) Mean mtDNA copy number in small (<0.3 μm) oocyte mitochondria. (H) Mean mtDNA copy number in large (>0.6 μm) oocyte mitochondria. Data are presented as mean ± SEM. * p<0.05, ** p<0.01 by Mann-Whitney U two-tailed test. ns, not significant.

Oocyte mitochondria were first separated into high ΔΨ_M_ and low ΔΨ_M_ subpopulations for young, aged, and NMN-supplemented aged samples (Figure 3C). In the high ΔΨ_M_ subpopulation, mitochondria from aged oocytes had a significantly elevated mtDNA copy number compared to those from young individuals (Figure 3D; mean = 0.303 ± 0.134 for young, 1.000 ± 0.282 for aged; n = 33 young, n = 33 aged; p = 0.0165). NMN supplementation of aged mice significantly reduced mtDNA copy number in this subpopulation relative to untreated aged mice, with values not significantly different from young mice (Figure 3D; n = 24 NMN-supplemented; p = 0.0056 for aged vs. NMN-supplemented). In the low ΔΨ_M_ subpopulation, mtDNA copy number did not vary significantly between young and aged oocytes, and NMN supplementation produced no significant difference from either age group (Figure 3E).

Mitochondrial subpopulations were also defined on the basis of size. Aged samples contained an increased proportion of large mitochondria relative to young samples (Figure 3F). In the small mitochondrial subpopulation (less than 0.3 μm), aged individuals had a significantly elevated average mtDNA copy number compared to young individuals (Figure 3G; mean = 0.229 ± 0.092 for young, 1.756 ± 0.728 for aged; n = 35 young, n = 45 aged; p = 0.0355). NMN supplementation of aged mice significantly reduced mtDNA copy number in small mitochondria relative to untreated aged individuals, with values not significantly different from young mice (Figure 3G; n = 24, mean = 0.21, SEM = 0.085, p = 0.040 vs. aged untreated). In the large mitochondrial subpopulation (greater than 0.6 μm), mtDNA copy number did not differ significantly between young and aged samples (Figure 3H; mean = 1.200 ± 0.389 for young, 0.788 ± 0.260 for aged; n = 35 young, n = 35 aged; p = 0.5424). NMN supplementation of aged individuals resulted in a mtDNA copy number in large mitochondria that was not significantly different from untreated aged mice but was significantly lower than young mice (Figure 3H; n = 24, mean = 0.21, SEM = 0.104, p = 0.017 vs. young). Taken together, these subpopulation findings reveal age- and NMN-associated differences in oocyte mitochondria that are not detectable at the aggregate level, and demonstrate that NMN exerts subpopulation-specific effects rather than acting uniformly across the mitochondrial pool.

### In cumulus cell mitochondria, mtDNA content and ΔΨ_M_ vary significantly with age

Cumulus granulosa cells (CCs) provide direct metabolic support to the developing oocyte through a network of gap junctions, supplying nutrients including pyruvate that oocyte mitochondria rely upon for energy production [28,10]. Given this metabolic interdependence, age-associated changes in CC mitochondrial function are likely to have downstream consequences for oocyte quality. CC mitochondria from young, aged, and NMN-supplemented aged mice were therefore assessed on the basis of size, ΔΨ_M_, and mtDNA copy number using the same FAMS and smPCR approach applied to oocytes.

At the aggregate level, CC mitochondria from aged individuals showed both a significant decrease in mtDNA copy number (Figure 4A; mean = 2.552 copies per mitochondrion in young [n = 67, SEM = 0.410] vs. 1.254 copies per mitochondrion in aged [n = 63, SEM = 0.302]; p = 0.0013) and a significant increase in ΔΨM (Figure 4B; 55.33% ΔΨ_M_-positive in young [n = 9, SEM = 1.968] vs. 66.67% in aged [n = 6, SEM = 2.769]; p = 0.0006). NMN supplementation in aged mice had a rejuvenative effect on both measures at the aggregate level. ΔΨ_M_ in NMN-supplemented aged mice was not significantly different from either the young or aged values, reflecting a partial reduction toward young levels (Figure 4B; n = 6, SEM = 2.762). mtDNA copy number showed a full rescue, with NMN-supplemented CC mitochondria having significantly higher mtDNA copy numbers than aged CC mitochondria and no significant difference from young CC mitochondria (Figure 4A; n = 37, SEM = 0.236).

**Figure 4.**
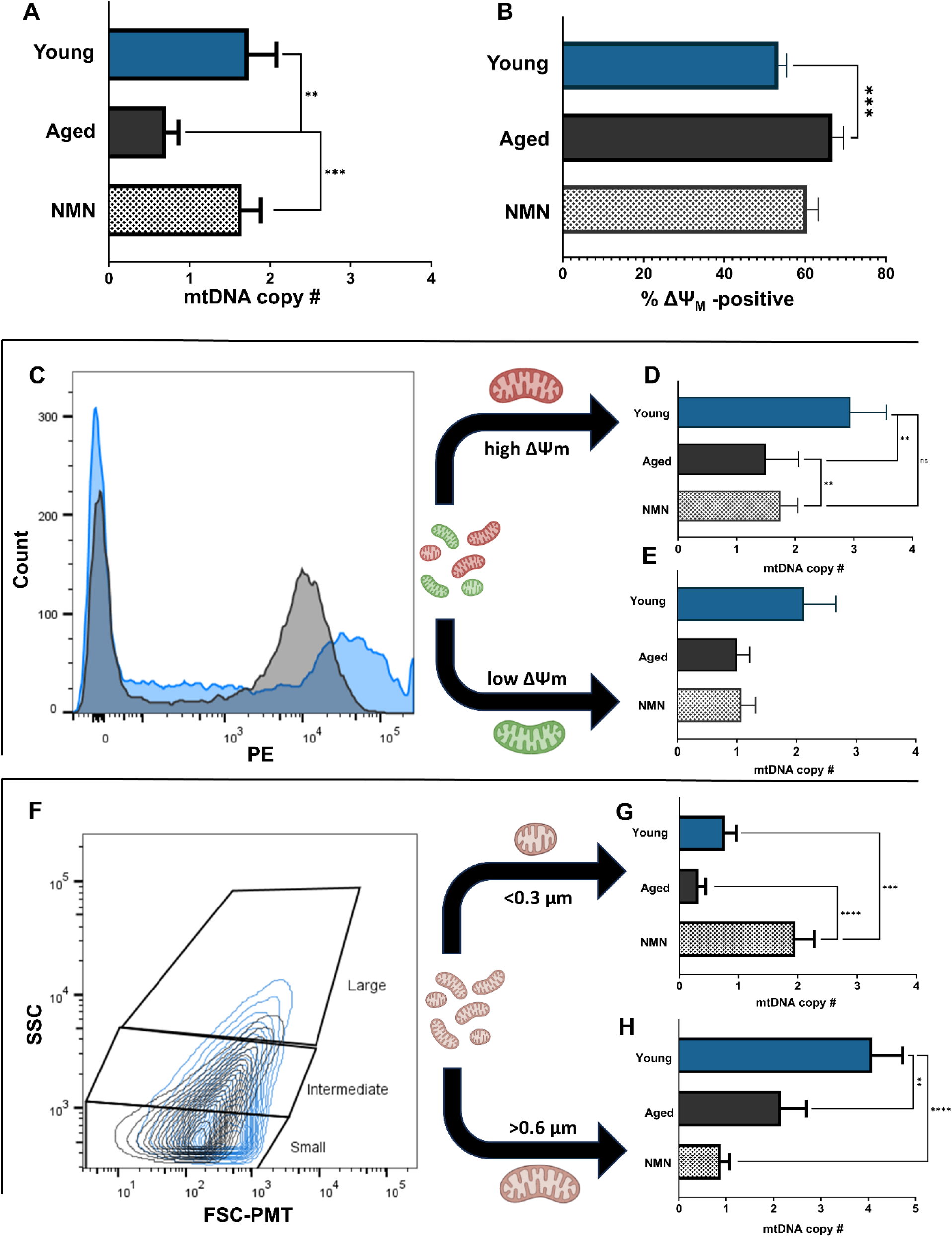
Age- and NMN-associated changes in mtDNA copy number and ΔΨM in cumulus cell mitochondrial subpopulations. Mitochondria isolated from cumulus granulosa cells of young (2-month-old, blue), aged (12-month-old, black), and NMN-supplemented aged (hatched gray) C57BL/6 female mice were analyzed by FAMS and assessed for mtDNA copy number by smPCR. (A) Mean mtDNA copy number per mitochondrion across the total cumulus cell mitochondrial population. (B) Percentage of ΔΨM-positive mitochondria across the total cumulus cell mitochondrial population. (C) Representative JC-1 PE channel histogram showing the distribution of ΔΨM-positive and ΔΨM-negative mitochondria in young (light blue) and aged (dark gray) cumulus cell samples, demonstrating the age-associated shift toward a higher proportion of ΔΨM-positive mitochondria. Curved arrows indicate separation into high and low ΔΨM subpopulations. (D) Mean mtDNA copy number in the high ΔΨM cumulus cell mitochondrial subpopulation. (E) Mean mtDNA copy number in the low ΔΨM cumulus cell mitochondrial subpopulation. (F) Representative FSC PMT vs. SSC contour plot showing the size distribution of cumulus cell mitochondria with small (<0.3 μm), intermediate (0.31–0.6 μm), and large (>0.6 μm) size gates indicated. Curved arrows indicate separation into small and large subpopulations. (G) Mean mtDNA copy number in small (<0.3 μm) cumulus cell mitochondria. (H) Mean mtDNA copy number in large (>0.6 μm) cumulus cell mitochondria. Data are presented as mean ± SEM. * p<0.05, ** p<0.01, *** p<0.001, **** p<0.0001 by Mann-Whitney U two-tailed test. ns, not significant.

Subpopulation analysis was then conducted to determine whether the aggregate differences were distributed uniformly across mitochondrial subtypes or concentrated within specific subpopulations. When CC mitochondria were divided on the basis of ΔΨ_M_, the aged sample contained a greater number of ΔΨ_M_-positive mitochondria, though their fluorescent peak was less intense than in young samples (Figure 4C). Aged ΔΨ_M_-positive mitochondria had a significantly reduced mtDNA copy number compared to young individuals (Figure 4D; mean = 2.943 ± 0.616 for young, 1.500 ± 0.559 for aged; n = 35 young, n = 32 aged; p = 0.0063). NMN supplementation significantly increased mtDNA copy number in this subpopulation relative to aged untreated mice, with values not significantly different from young CC mitochondria (Figure 4D; n = 32 aged, n = 24 NMN-supplemented; p = 0.0063 for aged vs. NMN-supplemented). In ΔΨ_M_-negative mitochondria, mtDNA copy number did not differ significantly between young and aged samples, and NMN supplementation produced no significant change (Figure 4E; mean = 2.125 ± 0.533 for young, 1.000 ± 0.213 for aged; n = 32 young, n = 31 aged; p = 0.0844; mean = 1.07 ± 0.238 for NMN-supplemented, n = 27). The age-associated reduction in mtDNA copy number across the total CC mitochondrial population was therefore driven predominantly by changes within the ΔΨ_M_-positive subpopulation.

When CC mitochondria were characterized on the basis of size, mtDNA copy number in small mitochondria did not change significantly with age (Figure 4G; mean = 0.774 ± 0.190 for young, 0.323 ± 0.117 for aged; n = 31 young, n = 31 aged; p = 0.0529). NMN supplementation of aged mice substantially increased mtDNA copy number in small CC mitochondria, with values significantly higher than both young and aged samples (mean = 1.958 ± 0.321; p = 0.001 vs. young, p < 0.001 vs. aged). In large CC mitochondria, mtDNA copy number was significantly reduced in aged samples compared to young samples (Figure 4H; mean = 4.083 ± 0.648 for young, 2.156 ± 0.541 for aged; n = 36 young, n = 32 aged; p = 0.002). NMN supplementation further reduced mtDNA copy number in large CC mitochondria, with values significantly lower than both aged and young samples (mean = 0.889 ± 0.180; p < 0.001 vs. young, p = 0.0703 vs. aged). These opposing effects of NMN on small and large CC mitochondrial subpopulations demonstrate that NMN supplementation does not universally restore CC mitochondria toward a youthful state, and that its effects on this cell type are subpopulation-specific. A summary of the direction and reversibility of all age- and NMN-associated changes across oocyte and cumulus cell mitochondrial subpopulations is provided in Figure 5.

**Figure 5.**
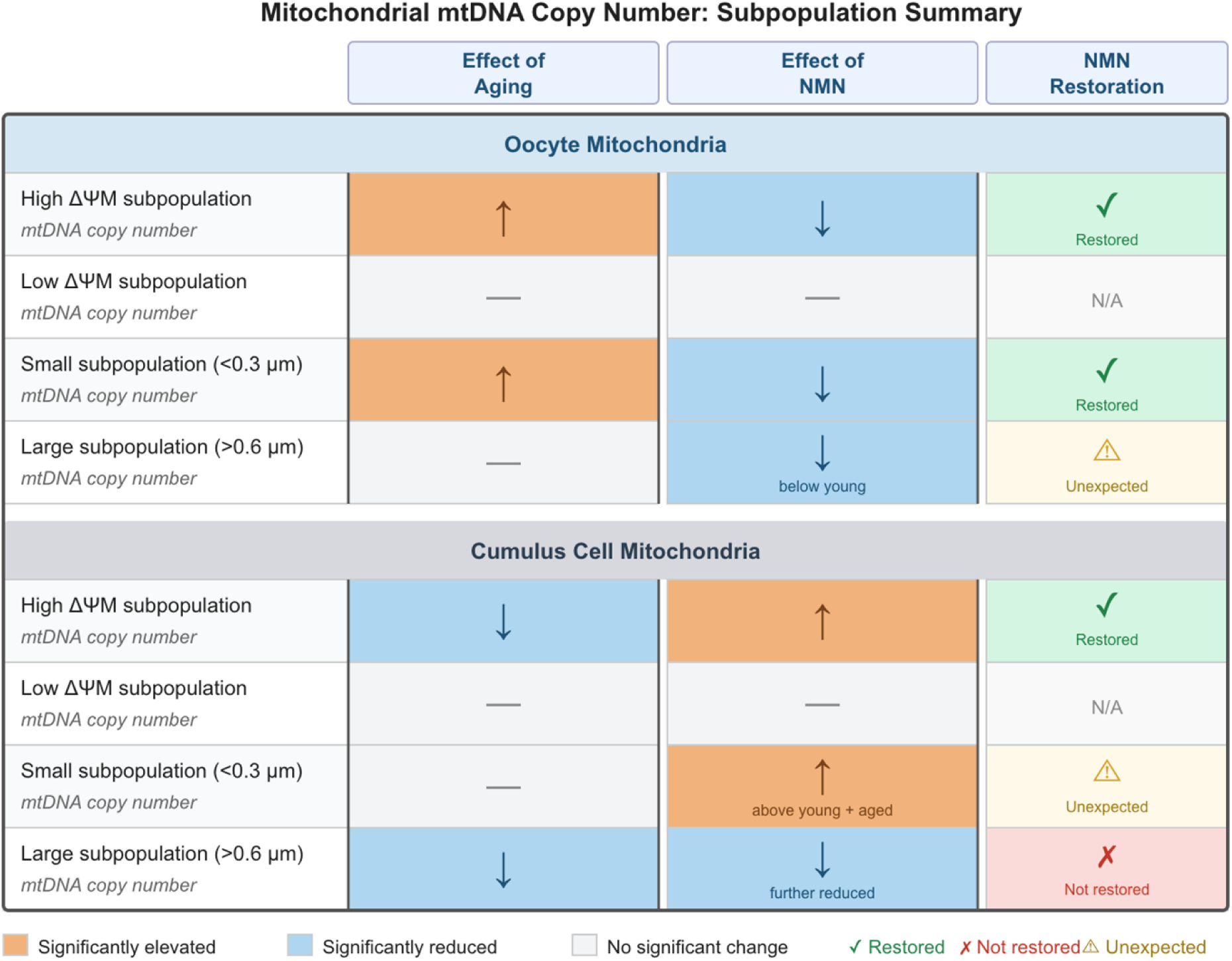
Summary of age- and NMN-associated changes in mtDNA copy number across mitochondrial subpopulations in oocytes and cumulus granulosa cells. Directional changes in mtDNA copy number are shown for each mitochondrial subpopulation identified by FAMS in oocytes (upper section) and cumulus granulosa cells (lower section). Subpopulations were defined on the basis of mitochondrial membrane potential (high ΔΨM and low ΔΨM) and size (small, <0.3 μm; large, >0.6 μm). The Effect of Aging column indicates the direction of significant change in mtDNA copy number between young (2-month-old) and aged (12-month-old) individuals. The Effect of NMN column indicates the direction of significant change in mtDNA copy number between untreated aged and NMN-supplemented aged individuals. The NMN Restoration column summarizes whether NMN supplementation restored the aged subpopulation phenotype toward young levels: ✓ (green) indicates restoration to values not significantly different from young; ✗ (red) indicates failure to restore; ⚠ (amber) indicates an unexpected effect in which NMN produced a significant change in a subpopulation that did not differ significantly between young and aged individuals, or drove values beyond young levels. Orange fill, significantly elevated; blue fill, significantly reduced; gray fill, no significant change. All comparisons by Mann-Whitney U two-tailed test, p<0.05.

## Discussion

Aging differentially affects distinct mitochondrial subpopulations in oocytes and cumulus granulosa cells in ways that aggregate approaches cannot detect, and the effects of NMN supplementation are similarly subpopulation-specific rather than uniform across the mitochondrial pool. A recurring and conceptually important theme across both cell types is the discordance between aggregate and subpopulation-level findings. In oocytes, neither mtDNA copy number nor ΔΨ_M_ differed significantly with age when the mitochondrial population was evaluated as a whole, which would previously have been interpreted as evidence that oocyte mitochondria are not meaningfully altered during reproductive aging. Subpopulation analysis revealed this conclusion to be incomplete: significant and opposing changes in mtDNA copy number were present within specific mitochondrial subpopulations, and these changes were obscured at the aggregate level precisely because they did not occur uniformly across the pool. In cumulus cells, significant differences were detectable at the aggregate level, yet subpopulation analysis further resolved these changes, demonstrating that the age-associated reduction in mtDNA copy number was driven predominantly by the ΔΨ_M_-positive subpopulation and that NMN supplementation acted differentially across size-defined subpopulations in ways the aggregate data could not capture. The implication is straightforward: bulk mitochondrial assays, including quantitative PCR-based mtDNA quantification and population-level membrane potential measurements, report on the average behavior of a heterogeneous organelle pool and may therefore systematically misrepresent the biology of specific functionally distinct subpopulations. Resolving individual mitochondrial subpopulations is not merely a technical refinement but an important consideration for accurately characterizing how aging affects the mitochondrial landscape of the ovarian follicle. The subpopulation-specific nature of these findings, together with the selective capacity of NMN to restore specific subpopulations, is summarized in Figure 5.

Within the high ΔΨ_M_ oocyte mitochondrial subpopulation, aged individuals displayed a significantly elevated mtDNA copy number relative to young individuals, a finding that is at first glance at odds with whole-cell studies reporting decreased oocyte mtDNA content with age [25–27]. However, these observations are not necessarily contradictory. It is plausible that a global age-associated reduction in total mitochondrial number drives the whole-cell decline in mtDNA detected by qPCR-based approaches, while within the surviving organelles, particularly those maintaining high ΔΨ_M_, mtDNA copy number is paradoxically elevated. Several non-mutually exclusive mechanisms could account for this elevation. Compensatory mtDNA replication may occur within functionally active mitochondria in response to an overall decline in mitochondrial mass, as has been proposed in other aging contexts [29]. Alternatively, impaired mitophagy clearance could result in the retention of mtDNA-rich, high ΔΨ_M_ mitochondria that would otherwise be targeted for degradation. This possibility is of particular interest given recent work from our laboratory demonstrating that PINK1/Parkin-mediated mitophagy is preferentially associated with high ΔΨ_M_ mitochondria and that mitophagy-targeted organelles carry significantly elevated mtDNA content [14]. A third possibility is that age-associated changes in mitochondrial fission dynamics lead to altered mtDNA distribution across the mitochondrial pool, with small mitochondria accumulating disproportionate mtDNA as a consequence of asymmetric fission events. Distinguishing between these mechanisms will require direct assessment of mitophagy flux and mtDNA replication rates at the single-organelle level and represents an important direction for future investigation.

The parallel elevation of mtDNA copy number in small oocyte mitochondria from aged individuals mirrors the pattern observed in the high ΔΨ_M_ subpopulation. Recent work from our laboratory has established that large oocyte mitochondria exhibit significantly higher ΔΨ_M_ than their smaller counterparts, with ΔΨ_M_ increasing progressively as a function of mitochondrial size (H. Sheehan and colleagues, unpublished observations), suggesting that the size- and ΔΨ_M_-based subpopulation findings reported here are likely to reflect overlapping biological phenomena rather than entirely independent processes. The convergence of elevated mtDNA across two independently defined subpopulations therefore strengthens the conclusion that age-associated mtDNA dysregulation in oocytes is subpopulation-specific and selectively concentrated in the more metabolically active organelles.

To assess whether aged mitochondrial subpopulations retain the capacity for change, NMN supplementation was used as a pharmacological probe of mitochondrial plasticity rather than as a candidate therapeutic intervention. The ability of NMN to alter specific mitochondrial subpopulations in aged individuals therefore reflects the reversibility of age-associated changes rather than simply the efficacy of NMN as a fertility treatment. NMN supplementation normalized mtDNA copy number in both the high ΔΨ_M_ and small oocyte mitochondrial subpopulations toward young levels, consistent with the broader rejuvenative effects of NMN on oocyte mitochondrial dynamics reported previously [9,18]. Notably, NMN did not significantly affect the low ΔΨ_M_ or large mitochondrial subpopulations in the same manner, indicating that its effects are not uniform across the mitochondrial pool. In large oocyte mitochondria specifically, NMN supplementation resulted in a mtDNA copy number that was significantly reduced compared to young individuals, though not significantly different from aged individuals. While this pattern does not fit a straightforward rejuvenation narrative, it may reflect NMN-driven promotion of mitochondrial fission in aged oocytes, which has been reported in prior studies of NMN’s effects on mitochondrial dynamics [9,24], and which would be expected to reduce the mtDNA content of large mitochondria by redistributing it into smaller daughter organelles. This interpretation is speculative and will require direct assessment of fission and fusion dynamics in NMN-supplemented aged oocytes to confirm.

The functional significance of the large mitochondrial subpopulation in oocytes may extend beyond bioenergetics. Recent work from our laboratory has demonstrated that large, high-ΔΨ_M_ mitochondria in mouse oocytes and preimplantation embryos are preferentially enriched for the trophectoderm-specification transcription factor Tead4, implicating this subpopulation in the first mammalian cell fate decision (H. Sheehan and colleagues, unpublished observations). Age-associated alterations within this subpopulation, and their partial but incomplete normalization by NMN, may therefore carry developmental consequences that extend into the preimplantation period and warrant investigation in future studies. Taken together, the subpopulation-specific nature of both the age-associated changes and the NMN response in oocyte mitochondria points to these individual subpopulations as more precise targets for future mechanistic studies of reproductive aging than the oocyte mitochondrial pool considered in aggregate.

The mitochondrial changes observed in oocytes with age and in response to NMN supplementation do not occur in isolation; given the direct metabolic interdependence between oocytes and their surrounding cumulus cells, alterations in CC mitochondrial subpopulations are likely to influence oocyte bioenergetics and developmental competence. Subpopulation analysis of CC mitochondria on the basis of size revealed a more complex pattern of age- and NMN-associated changes than those observed at the aggregate level, and one that does not conform to a uniform rejuvenation model. In small CC mitochondria, mtDNA copy number did not differ significantly between young and aged individuals, yet NMN supplementation of aged mice produced a striking increase in mtDNA copy number that was significantly elevated above both young and aged values. In large CC mitochondria, the pattern differed: aged individuals showed a significant reduction in mtDNA copy number relative to young individuals, and NMN supplementation further exacerbated this reduction, driving mtDNA copy number significantly below both the aged and young values. These opposing effects of NMN across the small and large CC mitochondrial subpopulations demonstrate that NMN does not act uniformly on the mitochondrial pool of cumulus cells, and that its effects are subpopulation-specific in ways that would be entirely obscured by aggregate analysis.

The mechanistic basis for these divergent NMN effects remains to be determined, and two non-mutually exclusive possibilities are worth considering. First, NMN-driven promotion of mitochondrial fission, which has been reported in aged ovarian cells [9,24], would be expected to fragment large mitochondria into smaller daughter organelles, redistributing mtDNA from the large to the small subpopulation. Under this model, the simultaneous reduction in large mitochondrial mtDNA and elevation in small mitochondrial mtDNA observed with NMN supplementation would reflect a single underlying process of fission-driven redistribution rather than two independent phenomena. Second, it is possible that distinct mitochondrial subpopulations differ in their sensitivity to NAD+ repletion, such that small and large CC mitochondria respond to NMN through different downstream pathways with opposing effects on mtDNA maintenance. These possibilities are not mutually exclusive and may act in concert; distinguishing between them will require direct assessment of mitochondrial fission and fusion dynamics alongside mtDNA replication rates in NMN-supplemented CC mitochondrial subpopulations. Regardless of mechanism, the finding that NMN does not restore all CC mitochondrial subpopulations uniformly toward a youthful state is an important qualification to the broader rejuvenation narrative and highlights the value of subpopulation-level analysis in evaluating the effects of candidate fertility-preserving interventions.

The observation that aged CCs harbor a greater proportion of ΔΨ_M_-positive mitochondria alongside reduced mtDNA copy number within this subpopulation warrants careful interpretation. Classically, mitochondrial depolarization has been considered a prerequisite for PINK1/Parkin-mediated mitophagy; however, recent work from our laboratory using FAMS has demonstrated that PINK1/Parkin colocalization is in fact predominantly associated with high ΔΨ_M_ mitochondria, with low ΔΨ_M_ mitochondria showing minimal PINK1/Parkin colocalization [14]. Under this revised framework, the accumulation of high ΔΨ_M_ mitochondria in aged CCs may reflect an impairment of mitophagy flux rather than enhanced mitochondrial fitness, in that mitochondria primed for mitophagy are not being efficiently cleared. The concurrent reduction in mtDNA copy number within this subpopulation, which diverges from the elevated mtDNA copy number observed in PINK1/Parkin-positive mitochondria in our prior work, further supports the idea that aged CC mitochondria exhibit a disrupted relationship between membrane potential, mtDNA maintenance, and mitophagy. This dissociation between membrane potential and mtDNA copy number in aged CCs is particularly notable when compared to the oocyte findings, where elevated ΔΨ_M_ and elevated mtDNA copy number co-occurred within the same subpopulation. That the two cell types show divergent subpopulation-level responses to aging despite being metabolically coupled suggests that the mechanisms driving mitochondrial dysfunction with age may be cell type-specific, even within the same follicular microenvironment. The partial restoration of both ΔΨ_M_ distribution and mtDNA copy number in NMN-supplemented CCs is consistent with NMN improving mitophagy efficiency in aged granulosa cells, as reported previously [18], and suggests that the mitophagy impairment observed in aged CCs is reversible. Direct application of the PINK1/Parkin FAMS assay described by Piasecki et al. [14] to aged and NMN-supplemented CC mitochondria would be a logical and informative extension of the present findings.

### Conclusions

Aging differentially affects distinct mitochondrial subpopulations in oocytes and cumulus granulosa cells in ways that are not detectable when mitochondria are assessed in aggregate, demonstrating that subpopulation-level analysis is necessary for an accurate characterization of mitochondrial dynamics during reproductive aging. The divergent subpopulation-level responses to aging observed between oocytes and cumulus cells, despite their metabolic interdependence, suggest that the mechanisms driving mitochondrial dysfunction with age are cell type-specific even within the same follicular microenvironment. The plasticity of aged mitochondria demonstrated here, particularly their capacity to respond selectively to NAD+ repletion, supports the hypothesis that age-associated mitochondrial dysfunction in the ovarian follicle is driven by the cellular environment rather than by irreversible intrinsic changes to the organelles themselves. Future studies directly assessing mitophagy flux, mtDNA replication rates, and fission and fusion dynamics within individual mitochondrial subpopulations will be essential to fully characterize the mechanisms identified here, and to determine whether targeted modulation of specific subpopulations represents a viable strategy for preserving or restoring fertility in aged individuals.

## List of abbreviations

FAMS: Fluorescence-Activated Mitochondria Sorting
mtDNA: mitochondrial DNA
ΔΨ_M_: mitochondrial membrane potential
NMN: nicotinamide mononucleotide
NAD+: nicotinamide Adenine Dinucleotide
COCs: cumulus-oocyte-complexes
PMSG: pregnant mare serum gonadotropin
hCG: human chorionic gonadotropin
MTG: MitoTracker Green
FSC: forward light scatter
SSC: side light scatter
smPCR: single molecule polymerase chain reaction
CCs: cumulus granulosa cells

## Declarations

### Data Availability Statement

The data presented in this study are included in the article and further inquiries can be directed to the corresponding author.

### Ethics Statement

All experiments described herein were reviewed and approved by the Institutional Animal Care and Use Committee at Northeastern University (Protocol #20-0315R).

### Author Contributions

D.C.W. and J.L.T. conceived and designed the project. A.J.P., and H.C.S. analyzed the data. H.C.S. designed all protocols. A.J.P. and H.C.S. wrote the manuscript and prepared the display items. H.C.S., A.J.P., and P.L.H. conducted experiments. All authors reviewed and approved the final manuscript.

### Funding

This work was supported by grants from the National Science Foundation (2227756 to J.L.T. and D.C.W.) and A.J.P. was supported by the National Science Foundation’s Graduate Research Fellowship Program (DGE-1938052).

### Competing Interests

D.C.W. and J.L.T. declare interest in intellectual property described in U.S. Patent 8,642,329, U.S. Patent 8,647,869, U.S. Patent 9,150,830, and U.S. Patent 10,525,086. Hannah Sheehan is the CEO and CSO of SauveBio, Inc. The remaining authors declare that the research was conducted in the absence of any commercial or financial relationships that could be construed as a potential conflict of interest.

### Consent for publication

Not Applicable

## Acknowledgements

A previous version of this manuscript was included in a dissertation titled “Characterization of subpopulations within the cellular and intracellular landscape of the aging mammalian ovary” submitted by Hannah Sheehan to the College of Science of Northeastern University in partial fulfillment of the requirements for the degree of Doctor of Philosophy.

